## Supplementary File 1 for "Bronchus-associated macrophages efficiently capture and present soluble inhaled antigens and are capable of local Th2 cell activation"

**Supplementary file 1**: Table of antibodies and reagents used for flow cytometry and microscopy

| Antigen/reagent | Fluorophores | Vendor | Clone |
| --- | --- | --- | --- |
| CCR7 | Biotin | Biolegend | 4B12 |
| CD103 | PerCP-Cy5.5 | Biolegend | 2E7 |
| CD11b | BV785 | Biolegend | M1/70 |
| CD11b | FITC | Caltag | M1/70.15 |
| CD11c | BV650, PE-Cy7 | Biolegend | N418 |
| CD16/32  (TruStain FcX) | Purified | Biolegend | 93 |
| CD169 | PE, eFluor660 | eBioscience | Ser-4 |
| CD19 | PE-Dazzle594 | Biolegend | 6D5 |
| CD19 | PE-Cy7 | BD | 1D3 |
| CD206 | Biotin, PE | Biolegend | C068C2 |
| CD206 | PE | eBioscience | MR6F3 |
| CD24 | BV421 | Biolegend | M1/69 |
| CD24 | APC | eBioscience | M1/69 |
| CD4 | APC-H7 | BD | GK1.5 |
| CD4 | Pacific Blue | Biolegend | RM4-5 |
| CD4 | PE-Cy7 | eBioscience | GK1.5 |
| CD45 | Alexa 700 | Biolegend | 30-F11 |
| CD45.1 | Pacific Blue, Alexa 700 | Biolegend | A20 |
| CD45.1 | PE | BD | A20 |
| CD45.2 | Alexa 700 | Biolegend | 104 |
| CD64 | PE, PE-Cy7 | Biolegend | X54-5/7.1 |
| CD80 | Biotin | UCSF MAb core | 16.10.A1 |
| CD86 | Biotin | UCSF MAb core | GL-1 |
| I-A^b^ | Biotin, FITC, Alexa 647 | Biolegend | KH74 |
| IFNγ | Alexa 488 | Biolegend | XMG1.2 |
| IL-13 | PE | eBioscience | eBio13A |
| L1CAM (CD171) | PE | Miltenyi Biotec | 555 |
| Ly6C | BV711 | Biolegend | HK1.4 |
| Ly6G | BV510, PE-Cy7 | Biolegend | 1A8 |
| MerTK | Biotin | R&D | polyclonal |
| MerTK | PE | Biolegend | 2B10C42 |
| Siglec-F | BV421, PE, PE-CF594 | BD | E50-2440 |
| Streptavidin | Qdot605 | Life Technologies |  |
| TCRβ | APC | Biolegend | H57-597 |
| Thy1.1 | PE-Cy7 | Biolegend | OX-7 |
| Viability dye | eFluor780 | eBioscience |  |
| Vα2 | PE | BD | B20.1 |
| Vβ5.1/.2 | FITC | BD | MR9-4 |
| Vβ5.1/.2 | PerCP-e710 | eBioscience | MR9-4 |
| Y-Ae | Biotin | eBioscience | eBioY-Ae |
