## Supplementary File 2 for "Bronchus-associated macrophages efficiently capture and present soluble inhaled antigens and are capable of local Th2 cell activation"

**Supplementary file 2:** Table of emission filters for 2-photon microscopy

| Emission/Fluorophore | Filter | Vendor |
| --- | --- | --- |
| Second harmonic  (half of incident laser wavelength) | 450 short-pass  460/60  469/35  470/40  510/20  539/30 | Chroma  Semrock  Semrock  Chroma  Chroma  Semrock |
| CFP | 469/35  472/30 | Semrock  Semrock |
| GFP | 525/50  510/20  510/20 | Chroma  Chroma  Semrock |
| YFP | 525/50  525/40  535/30  539/30 | Chroma  Semrock  Chroma  Semrock |
| PE | 588/45  610/75 | Zeiss  Chroma |
| tdTomato | 588/45  605/70  610/75  625/30 | Zeiss  Chroma  Chroma  Chroma |
| Tetramethylrhodamine  (TRITC, TAMRA) | 610/75  625/30 | Chroma  Chroma |
| Texas Red | 605/70  610/75  625/30 | Chroma  Chroma  Chroma |
| Alexa 647 | 670/50  690/50 | Chroma  Zeiss |
| eFluor 660 | 690/50 | Zeiss |
| Sky Blue beads | 690/50 | Zeiss |
