## Supplementary File 3 for "Bronchus-associated macrophages efficiently capture and present soluble inhaled antigens and are capable of local Th2 cell activation"

**Supplementary file 3:** Table of excitation wavelengths for 2-photon microscopy

| Fluorophore | Excitation Wavelengths  (nm) |
| --- | --- |
| CFP | 855-890 |
| GFP | 855-940 |
| YFP | 890-1040 |
| PE | 1040-1140 |
| tdTomato | 890-1040 |
| Tetramethylrhodamine  (TRITC, TAMRA) | 870-890 |
| Texas Red | 860-1020 |
| Alexa 647 | 1100-1200 |
| eFluor 660 | 1140 |
| Sky Blue beads | 850 |
